# Evidence for peripheral and central actions of codeine to dysregulate deglutition in the anesthetized cat

**DOI:** 10.1101/2023.11.07.566014

**Authors:** Donald C. Bolser, Tabitha Y. Shen, M. Nicholas Musselwhite, Melanie J. Rose, John A. Hayes, Teresa G. Pitts

## Abstract

Systemic administration of opioids has been associated with aspiration and swallow dysfunction in humans. We speculated that systemic administration of codeine would induce dysfunctional swallowing and that this effect would have a peripheral component. Experiments were conducted in spontaneously breathing, anesthetized cats. The animals were tracheotomized and electromyogram electrodes were placed in upper airway and chest wall respiratory muscles for recording swallow related motor activity. The animals were allocated into three groups: vagal intact (VI), cervical vagotomy (CVx), and supra-nodose ganglion vagotomy (SNGx). A dose response to intravenous codeine was performed in each animal. Swallowing was elicited by injection of 3 ml of water into the oropharynx. The number of swallows after vehicle was significantly higher in the VI group than in SNGx. Codeine had no significant effect on the number of swallows induced by water in any of the groups. However, the magnitudes of water swallow-related EMGs of the thyropharyngeus muscle were significantly increased in the VI and CVx groups by 2-4 fold in a dose-related manner. In the CVx group, the geniohyoid muscle EMG during water swallows was significantly increased. There was a significant codeine dose-related increase in spontaneous swallowing in each group. The spontaneous swallow number at the 10 mg/kg dose of codeine was significantly larger in the CVx than that in the SNGx groups. During water-evoked swallows, intravenous codeine increased upper airway motor drive in a dose-related manner, consistent with dysregulation. The appearance of spontaneous swallowing in response to codeine in all groups supports a central action of this drug on the swallow pattern generator and also is consistent with dysregulation. At the highest dose of codeine, the reduced spontaneous swallow number in the SNGx group relative to CVx is consistent with a peripheral excitatory action of codeine either on pharyngeal/laryngeal receptors or in the nodose ganglion itself.

## INTRODUCTION

Swallow is a critically important behavior for protecting the airways from aspiration. Swallowing is produced by a complex neural circuit that is functionally identified as the swallow pattern generator. This circuit is located in the brainstem, but is subject to significant modulation by suprapontine pathways(1). Further, swallowing also is modulated by peripheral sensory afferents in the lungs and airway(2, 3), chest wall(2), mouth, pharynx, and esophagus(1, 4-6) and these afferents travel in the vagus, trigeminal, or hypoglossal nerves.

Administration of opioids, including morphine and remifentanyl is associated with significant dysphagia and aspiration syndromes (7-11). The site(s) of opioid actions on swallowing are not fully understood. It is well known that systemic administration of opioids can activate pulmonary vagal afferents resulting in perturbation of laryngeal function(12, 13). Further, microinjection of opioids into the nodose ganglion can induce effects that are similar to activation of peripheral vagal afferents(14). Savilampi *et al* (10) found in a study of a small number of subjects that subjective metrics of swallow difficulty and objective metrics of esopho-gastric junction activity during swallowing were not affected by pretreatment with a peripherally-restricted opioid antagonist. The results of this study were consistent with a peripheral action of opioids to influence swallowing in these subjects.

Codeine is one of the most commonly abused opioid drugs worldwide and is among the most common pharmaceutical opioids found in fatal poisonings (15, 16). Systemic administration of this drug is known to stimulate vagal C-fibers to influence breathing(17). Further, codeine is well known to alter breathing and inhibit cough via central action(18-21). The influence of this opioid on swallowing is unknown. We speculated that intravenous codeine would have suppressive effects on swallow frequency and upper airway muscle EMG magnitudes and that some of these effects would be mediated by vagal afferents.

## METHODS

Experiments were performed on 19 spontaneously breathing adult male cats with an average age of 1.05 years (SD = 0.14 years) and weight of 4.95 kg (SD = 0.53 kg). These cats were purchased from Marshall BioResources (North Rose, NY) and pair-housed in the University of Florida Animal Care Services on a 12-h light/12-h dark cycle with food and water ad libitum. The protocol was approved by the University of Florida Institutional Animal Care and Use Committee (IACUC) and all procedures were compliant with the Guide for the Care and Use of Laboratory Animals. Anesthesia was initially induced with sevoflurane (4.5%), then the animals were weaned onto sodium pentobarbital (Lundbeck, Inc., Deerfield, IL, 25 mg/kg, i.v.). Based on forelimb withdrawal reflex and jaw tone, supplemental doses were given as needed in 0.1-0.3 mL i.v. boluses. A single dose of atropine sulfate (0.054 mg/kg, i.v., Patterson Veterinary Supply, Inc., Devens, MA) was given at the beginning of surgery to minimize airway secretions. The femoral vein and artery were cannulated to deliver drugs and record blood pressure, respectively. Arterial blood pressure and end-tidal CO_2_ (ETCO_2_, (4–4.5%; Datax Engstrom; Datax Ohmeda, Inc; Madison, WI)) were continuously recorded, and arterial blood gas samples (i-STAT1; Abaxis; Union City, CA) were obtained every hour. Body temperature was maintained at 37.5 ± 0.5 °C with a TC-1000 homeothermic pad and rectal temperature probe (CWE Inc.; United States). Esophageal pressure was measured via a small balloon inserted through the mouth into the midthoracic esophagus. The cats were euthanized by an overdose of pentobarbital (iv) followed by 3 mL of a saturated potassium chloride solution (iv) (Thermo Fisher Scientific, Waltham, MA).

Swallow was induced by introducing 3 ccs of water into the pharynx via a 1-inch long, thin polyethylene catheter (outer diameter 0.5–1.0 mm) placed rostral to the faucial pillars and attached to a 5 cc syringe. All water trials for each animal were performed by the same researcher to maintain stimulus consistency. All stimuli were presented three times, separated by a minimum inter-stimulus interval of one minute.

Animals were tracheostomized and a bifurcated tracheostomy tube was placed to permit simultaneous end-tidal gas recording and manipulation of inspired gases. End-tidal gas was measured via a GEMINI O2 & CO2 Monitor (CWE Inc., Ardmore, PA). Electromyograms (EMGs) were recorded using bipolar fine wire electrodes placed intramuscularly to record respiratory related behaviors using the method of Basmajian and Stecko(22). EMGs from the sternocostal diaphragm (dorsal to the xiphoid process), bilateral parasternal, bilateral internal abdominal oblique, thyroarytenoid, posterior cricoarytenoid (PCA), geniohyoid, mylohyoid, thyropharyngeus and the upper esophageal sphincter were recorded to classify respiratory behaviors and to evaluate breathing, cough, and swallowing. EMG signals were amplified using Grass P511 amplifiers and band-pass filtered (300-10,000 Hz) and subsequently digitized with a Power 1401 mk II and Spike 2 version 9 (Cambridge Electronic Design; United Kingdom). EMG channels were rectified and smoothed with a 50 ms time constant prior to analysis. EMG amplitude measures were normalized to the largest swallow or breath and are presented as % of maximum. EMG signals were sampled at 25,000 Hz and pressure signals at 500 Hz.

Surgical placement of EMGs proceeded as follows: the digastric muscles were blunt dissected away from the surface of the mylohyoid and electrodes were placed medially in the left mylohyoid. A small horizontal incision was made at the rostral end of the right mylohyoid followed by an incision down the midline for approximately 5 mm to reveal the geniohyoid muscle. Electrodes were placed 1 cm from the caudal insertion of the geniohyoid muscle. The thyroarytenoid muscle electrodes were inserted through the cricothyroid window into the anterior portion of the vocal folds, which were visually inspected post-mortem. Minor rotation of the larynx and pharynx counterclockwise revealed the superior laryngeal nerve, which facilitated placement of the thyropharyngeus muscle electrodes. The thyropharyngeus is a fan shaped muscle with the smallest portion attached to the thyroid cartilage; electrodes were placed in the ventral, caudal portion of the muscle overlaying thyroid cartilage within 5 mm of the rostral insertion of the muscle. To place electrodes within the cricopharyngeus muscle, the larynx and pharynx were rotated counterclockwise to reveal the posterior aspect of the larynx. The edge of the cricoid cartilage was located by palpation and electrodes were placed in the cricopharyngeus muscle just cranial to the edge of this structure. Thyrohyoid muscle electrodes were inserted approximately 5 mm rostral to the attachment to the thyroid cartilage; those for the parasternal muscle were placed in the third intercostal space, just adjacent to the sternum, and the costal diaphragm EMGs were placed transcutaneously just under the xiphoid process. The positions of all electrodes were confirmed by visual inspection (following electrode placement and post-mortem) and by EMG activity patterns during breathing and swallow, as we have previously published (Pitts et al., 2013, 2015b, 2018; Spearman et al., 2014).

Animals were segregated into three groups: vagal intact (n=7), vagotomized at the mid-cervical level bilaterally (n=6), and vagotomized above the nodose ganglion bilaterally (supranodose vagotomy, n=6). All protocols were initiated at least 1-hour post vagotomy. Two to five water swallow trails were conducted prior to administration of vehicle (0.9% saline) for the purpose of scaling amplification of EMG magnitudes and assessment of swallow excitability. After administration of vehicle and each dose of codeine (0.1-10 mg/kg, iv, Sigma Aldrich Chemical Co. St. Louis, MO), 2-5 water swallow trials were conducted beginning at least one minute after each dose and separated by approximately 1 minute. All doses of codeine phosphate were calculated as their free base.

EMGs were normalized to the median amplitude recorded for that specific muscle during swallows elicited in the vehicle period. Data were then aggregated, resulting in an average for each measured parameter per vehicle or dose of codeine. All data are reported as a mean and standard error unless otherwise stated. Normality of the data was tested using the Kolmogorov–Smirnov normality test. For statistical analysis of more than two groups, we used a repeated-measures ANOVA (RM-ANOVA) or an ordinary ANOVA with a Student–Newman–Keuls post-test for normally distributed data. If data did not pass the normality test, then group data were tested using the Kruskal–Wallis ANOVA with a Dunn’s post-test. Differences between swallow number pre- and post-cervical vagotomy and supranodose section were assessed with the Mann-Whitney test and a t-test with Welch correction, respectively. Alpha was set to 0.05.

## RESULTS

In the CVx and SNGx groups, pre-nerve section swallow trials were conducted for comparison to trials conducted 10-15 min later. In the CVx group, water swallow number was decreased after vagotomy from 2.0±0.38 to 1.1±0.29 (p<0.04). In the SNGx group, water swallow number was decreased from 4.0+0.97 to 0.75+0.46 (p<0.05). There was no significant difference between the pre-nerve section swallow numbers in the CVx and SNGx groups (p<0.07) although these values approached statistical significance. Further, there was no difference between the post section water swallow values for the CVx and SNGx groups (p<0.17).

Codeine induced spontaneous swallowing in all groups, especially at the highest dose (10 mg/kg, iv, Fig. 1). The number of swallows varied between groups. At the 3.0 mg/kg dose of codeine, the CVx swallow number (8.6±3.8) was significantly higher than that of either the VI (p<0.01, 1.0±3.6) or SNGx (p<0.05, 1.0±0.77) groups (Fig. 2). At the 10.0 mg/kg dose of codeine, the CVx swallow number (12.7±2.3) was significantly higher than that of either the VI (p<0.05, 5.5±1.9) or SNGx (p<0.01, 4.0±1.0) groups (Fig. 2).

**Figure 1.**
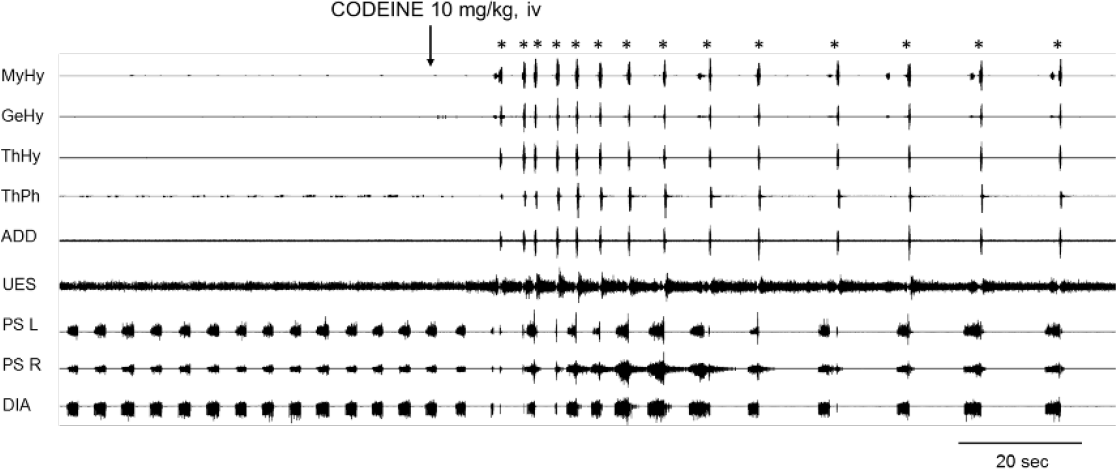
Intravenous codeine elicits spontaneous swallowing. Bolus administration of codeine (10 mg/kg, iv) transiently perturbed breathing and elicited multiple spontaneous swallows. MyHy-mylohyoid EMG, GeHy-geniohyoid EMG, ThHy-thyrohyoid EMG, ThPh-thyropharyngeus EMG, ADD-thyroarytenoid EMG, UES-upper esophageal sphincter EMG, PS L-left parasternal EMG, PS R-right parasternal EMG, DIA-diaphragm EMG. Swallows are labeled with asterisks.

**Figure 2.**
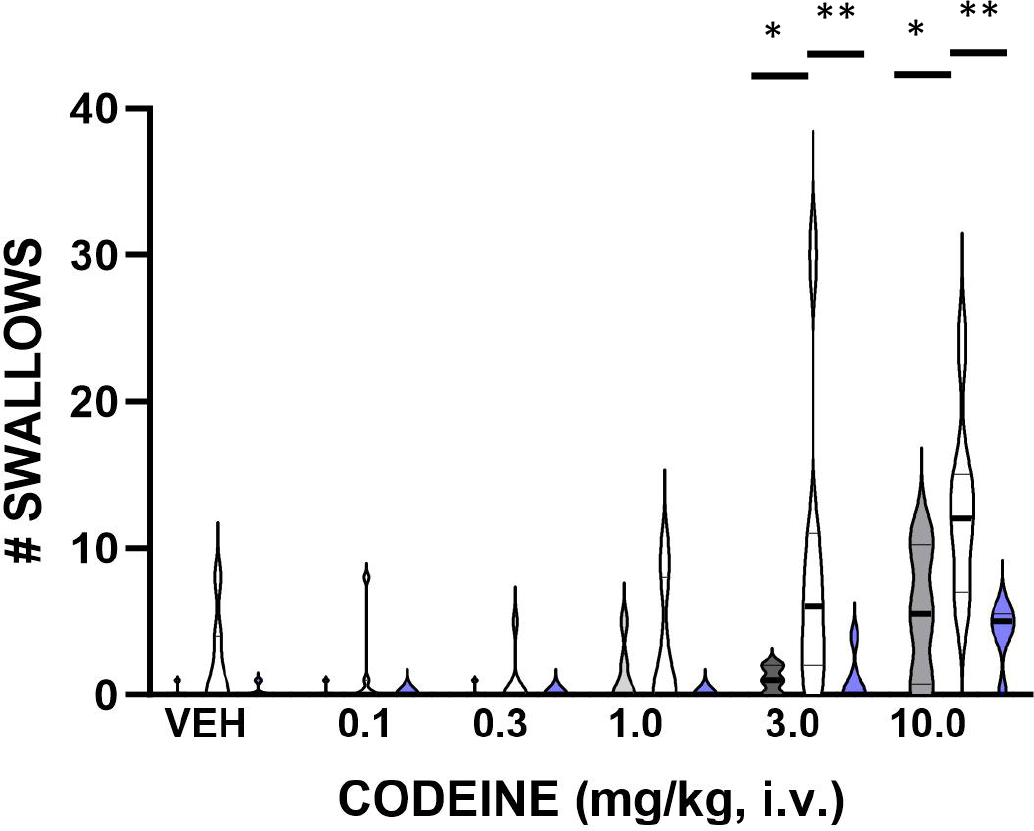
Violin plot of swallow number in response to increasing cumulative doses of codeine in VI, CVx, and SNGx groups. Codeine induced swallowing in all three groups but cervical vagotomized animals (n=7) exhibited significantly higher swallow numbers than vagal intact (n=5) or supranodose vagotomized animals (n=6). Grey-VI, open-CVx, blue-SNGx. *p<0.05, **p<0.01 relative to the CVx group.

There were no significant increases in swallow number after water trials in any of the three groups during the codeine dose response protocol. During water swallows, there were no statistically significant changes in upper airway EMG magnitudes in VI animals in response to administration of codeine (geniohyoid p<0.66; mylohyoid p<0.4; thyrohyoid p<0.94; thyropharyngeus p<0.48; thyroarytenoid p<0.95). In the SNGx group, there were very few swallows during water challenge trials so upper airway muscle EMGs were not analyzed. Cervical vagotomy did not alter the magnitudes of upper airway EMGs relative to their pre-vagotomy values (geniohyoid p<0.23; mylohyoid p<0.82; thyrohyoid p<0.31; thyropharyngeus p<0.17), although there were large variances in these values after vagotomy (geniohyoid, pre CVx 88±12, post CVx 106±4.5; mylohyoid, pre CVx 145±56, post CVx 106±3.9; thyrohyoid, pre CVx 126±22, post CVx 110±9.6; thyropharyngeus, pre CVx 151±33, post CVx 104±3.1).

In CVx animals, the EMG magnitudes of water swallows significantly increased after administration of codeine in the geniohyoid, mylohyoid, thyrohyoid and thyropharyngeus muscles. Given that recurrent laryngeal nerve motor axons were sectioned by vagotomy, thyroarytenoid EMG magnitudes were not analyzed in the CVx group. During individual swallows, upper airway muscle EMGs in CVx animals could be increased by over an order of magnitude (Fig. 3). The changes in magnitude were most pronounced at the 10 mg/kg dose of codeine (Fig. 3), although increases in EMG magnitude could be observed at lower doses in response to codeine. Geniohyoid swallow EMG magnitudes after 10 mg/kg codeine (p<0.01) were significantly larger than those in the vehicle period. However, there was no difference between the geniohyoid water swallow magnitudes after 10 mg/kg codeine in the VI and CVx groups (p<0.14). The magnitude of mylohyoid water swallow EMGs after 10 mg/kg codeine (p<0.05) were significantly larger than those in the vehicle period. The mylohyoid water swallow EMG magnitudes in the CVx group were significantly larger than those in the VI group at the 3.0 mg/kg (p<0.01) and 10.0 mg/kg (p<0.05) doses of codeine (Fig. 3). The magnitude of thyrohyoid water swallow EMGs after 10 mg/kg codeine (p<0.03) were significantly larger than those in the vehicle period. Thyrohyoid EMG magnitudes during water swallows in the CVx group were larger at the 3.0 (p<0.01) and 10.0 (p<0.001) doses of codeine than those in the vehicle period. There was no significant difference between the magnitudes of thyrohyoid EMGs during water swallow between the VI and CVx groups at any dose of codeine. Codeine induced significant increases in the thyropharyngeus EMG during water swallow relative to vehicle at the 3.0 mg/kg (p<0.01) and 10.0 mg/kg doses (p<0.001). There was no significant difference between the magnitudes of thyropharyngeus EMGs during water swallow between the VI and CVx groups at any dose of codeine.

**Figure 3.**
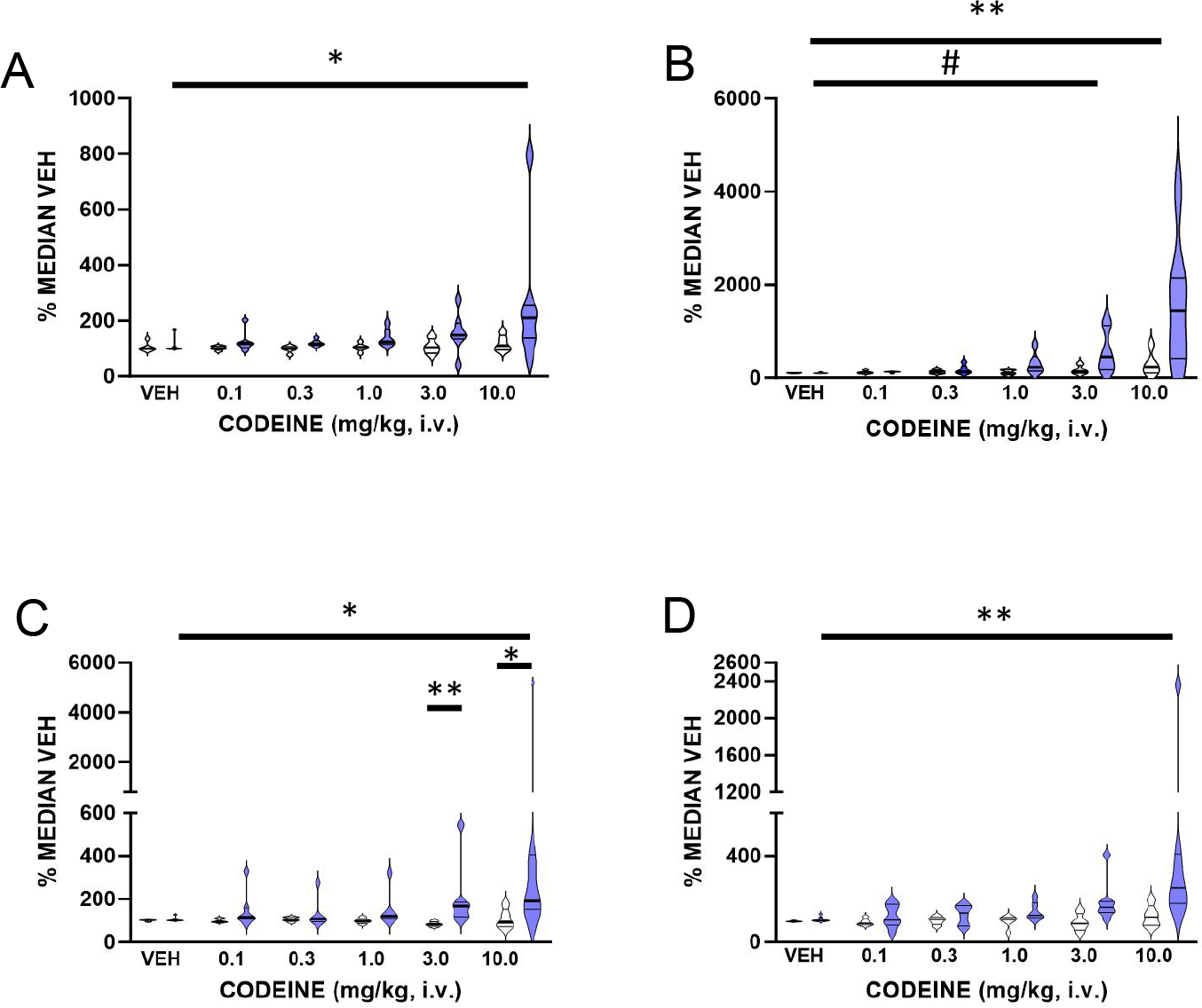
Influence of codeine on thyrohyoid (A), thyropharyngeus (B), mylohyoid (C), and geniohyoid (D) muscle EMG magnitudes during water swallows in VI and CVx animals. Codeine significantly enhanced EMG magnitudes in all four muscles in CVx animals. EMG data from SNGx animals not shown because there were few swallows in response to water in that group. VI (n=5 animals, CVx (n=7 animals). * p<0.05, ***p<0.03, **p<0.01, # p<0.001.

## DISCUSSION

The major findings of this study were that intravenous codeine induced spontaneous swallowing in a dose-dependent manner in vagal intact and vagally denervated cats. This drug did not increase the frequency of water-induced swallows in any group. During water swallows, codeine increased the EMG magnitudes of selected upper airway muscles (geniohyoid, mylohyoid, thyrohyoid and thyropharyngeus) only in animals with cervical vagotomy.

To our knowledge, this is the first report that the mu-opioid drug, codeine, alters swallow function in an animal model. In humans, the mu-opioid agonist, remifentanil, altered pharyngeal function and esophageal motility during swallowing (7-11, 23). These effects included frank aspiration (8), reductions in pharyngeal bolus movement (9), altered upper esophageal sphincter function (7), and a reduction in the strength of pharyngeal contractions (9). Similar effects were observed with morphine (9, 24). The effects of remifentanil on pharyngeal swallowing were not attenuated following administration of a peripherally acting derivative of the opioid antagonist naltrexone (7), suggesting a central action of the opioid. We are not aware of any report in the literature that codeine alters swallow function on the human. We note that we observed enhancement of pharyngeal swallow at doses of this drug (3 and 10mg/kg. iv) that were frankly respiratory depressant in this model (data not shown). In the human, prescription doses of codeine typically do not result in respiratory depression. It is possible that codeine in the human would have significant effects on swallow function at higher doses.

We cannot attribute these effects of codeine solely to its actions at mu-opioid receptors given that it is a nonspecific drug. Codeine binds to an allosteric site on the central nicotinic receptor and enhances cholinergic transmission by a mechanism that is not sensitive to the alpha-7 nicotinic receptor antagonist methyllycaconitine (25-27). Therefore, the excitatory effects on pharyngeal swallowing observed in this study may be specific to codeine and not extendable to mu-opioid drugs in general.

Codeine induced swallowing in the absence of a peripheral stimulus, and swallow number was higher in vagotomized relative to vagal intact or SNGx animals. This supports several different conclusions. First, codeine can induce swallowing in animals that have vagal axons and/or vagal axons plus ganglia sectioned, indicating that the swallow-promoting actions of this drug do not require sensory feedback from vagal sources to occur. Second, this vagal sensory feedback blunted the swallow-inducing effects of codeine. Third, the differences between cervical vagotomy and section of the vagal trunk rostral to the nodose ganglion support the concept that ganglion cells and/or superior laryngeal nerve afferent pathways had a role in blunting the actions of codeine. The superior and inferior laryngeal nerves enter the vagal ganglion complex and this anatomical site is rostral to the typical location of cervical vagotomy. This afferent pathway was intact in our CVx animals, but was eliminated in the SNGx group.

While codeine is well-known to have central actions (18, 28), it and other opioids also alter the excitability of thoracic vagal and perhaps laryngeal afferents (13, 17, 29). The extent to which the swallow-inducing actions of codeine were due to solely central effects of this drug that were modified by spontaneous discharge of vagal sensory pathways or were due to both central and peripheral actions is unknown.

Codeine had differential actions on swallowing when actuated by increasing doses of the drug itself or a water stimulus. Spontaneous swallow number increased in a dose-dependent manner in response to the drug, especially in CVx animals, yet the swallow number did not increase when the behavior was elicited by water. First, codeine may have acted to decrease the threshold for actuation of swallow, presumably at the central level. Second, codeine may have acted on the swallow central pattern generator for swallowing if it is subordinate to a distinct threshold mechanism. Third, this drug may have increased the excitability of other circuits that stimulate swallowing through disinhibition. For example, Pitts and coworkers have shown that spinal pathways can have significant excitatory effects on swallowing (30). Our approach cannot separate between these three hypotheses and they are not mutually exclusive.

These findings also support the concept that swallow frequency can be controlled differently than motor drive to upper airway muscles. In response to water stimuli, swallow number did not change but the magnitudes of upper airway muscle EMGs significantly increased, especially in CVx animals. As noted above, enhancement of motor drive to selected upper airway muscles during swallowing can be observed following cervical spinal injury (30). The extent to which this mechanism can be actuated by drugs in the absence of spinal injury is unknown.

We conducted these experiments solely in male animals. Recently, Huff et al (2) showed in a mouse model that there were no differences in swallow number or upper airway muscle EMG magnitudes between sexes under control conditions in vagal intact animals. However, male mice had 30-40% higher motor drive to geniohyoid and mylohyoid muscles after vagotomy. In our study, cervical vagotomy did not significantly alter motor drive to these two muscle groups or the thyrohyoid or thyropharyngeus muscles during water swallow but post vagotomy magnitudes were highly variable. The extent to which vagotomy alters upper airway muscle motor drive in female cats is unknown.

Bautista and co-workers have shown that swallowing can occur during autoresuscitation (31) and this behavior has been proposed to be important in the autoresuscitation process (32). It is likely that the role of swallowing in autoresuscitation is one of oropharyngeal clearance. Motor activation of pharyngeal muscles to move material out of the upper airway is an essential component of this behavior. Given that codeine initiated spontaneous swallowing but did not increase swallow number after water swallows, the action of this drug may be on neural elements that participate in non-ingestive functions of swallow control, such as autoresuscitation.

There were very large magnitude water swallows in our study for some upper airway muscle EMGs. This effect was magnified somewhat by the fact that vagotomy reduced EMG magnitudes for the thyropharyngeus in four animals. Although this effect was not statistically significant it did alter normalization. However, the largest water swallows in these animals were still several hundreds of percent larger than pre-vagotomy levels. As such, we chose not to remove these swallows from the dataset as outliers. It is our view that others who may choose to repeat our work be informed regarding the large range of swallow EMG magnitudes that can be observed after administration of codeine.

## Notes

### Competing Interest Statement

The authors have declared no competing interest.

